# How will it look like? Human posterior parietal and dorsal premotor cortex encode the visual properties of an upcoming action

**DOI:** 10.1101/322925

**Authors:** Artur Pilacinski, Melanie Wallscheid, Axel Lindner

## Abstract

Behavioral studies show that motor actions are planned by adapting motor programs to produce desired visual consequences. Does this mean that the brain plans these visual consequences independent of the motor actions required to obtain them? Here we addressed this question by investigating planning-related fMRI activity in human posterior parietal (PPC) and dorsal premotor (PMd) cortex. By manipulating visual movement of a virtual end-effector controlled via button presses we could dissociate motor actions from their sensory outcome. A clear representation of the visual consequences was visible in both PPC and PMd activity during early planning stages. Our findings suggest that in both PPC and PMd action plans are initially represented on the basis of the desired sensory outcomes while later activity shifts towards representing motor programs.

## INTRODUCTION

Reaches are realized through complex movements of individual joints, even though the resulting hand trajectories look surprisingly straight and have simple velocity profiles (Morasso, 1981). This suggested that the central nervous system aims at producing desired visual actions and adapts motor plans accordingly. In fact, Wolpert and colleagues (1995) confirmed that notion by an experiment, in which a mismatch between the actual hand position and the visual feedback thereof was manipulated. Despite this mismatch, subjects reached along visually straight trajectories. Similarly, related work by Mechsner and others (2001) revealed that subjects automatically preferred movements that led to visually symmetrical action-effects, even if the actual actions needed to generate these effects were not symmetrical. Several other lines of research have further suggested that movement plans indeed aim first at producing desired sensory outcomes (Janczyk et al 2014; Kunde, 2003; Shin et al. 2010; Kühn and Brass, 2010; Hommel, et al., 2001; Elsner et al., 2002; Wolpert and Ghahramani 2000). One may hypothesise that if these sensory representations are part of the action planning processes, then we should also be able to delineate neural substrates that contain them. Kühn and colleagues (2011) provided a valuable piece of evidence to support this idea, demonstrating that preparing hand vs. face actions also increases activity in visual areas related to the perception of body parts vs. faces, respectively (compare also to: Kühn et al., 2010). However, only little evidence directly demonstrates prospective representations of sensory action outcomes in the cortical areas engaged in action planning. It remains then an open question how the sensory outcomes become incorporated as part of action plans and what neural substrates are responsible for this.

One candidate brain area that could sub-serve such function in humans is the posterior parietal cortex (PPC). As repeatedly demonstrated, it plays a crucial role in forming movement intentions (Desmurget et al., 2009) and visual motor-imagery (Crammond, 1997; Sirigu et al., 1996). Importantly, the medial portion of human PPC has been considered a main substrate for reach planning (e.g. Lindner et al., 2010), possibly constituting the human homologue of macaque parietal reach region (PRR), as shown Connoly et al. (2003). Thereby, the PRR and its putative human homologue represent reach targets and effectors in a common visual reference frame (Andersen and Buneo, 2002; Buneo et al., 2002; Heed et al., 2011; Medendorp et al., 2008). In addition, other studies suggest that PPC/PRR processes also more complex aspects of upcoming movement, such as trajectory (Hauschild et al., 2012; Torres et al., 2013, Aflalo et al., 2015). Most direct evidence for PPC prospectively encoding the expected sensory consequences during action planning comes from a recent electrophysiological study in monkeys by Kuang et al. (2015). They combined anti-reach and prism adaptation paradigms in order to tease apart the motor and the visual properties of a reach. By using reversing prisms they were able to separate the visually perceived target location from the physical reach endpoint. Moreover, by using anti-reaches (reaches away from the sensory cue) they also could separate retrospective representations of the visible target cue from prospective planning. This manipulation allowed them to demonstrate, in one monkey, that at least some PPC neurons do encode the predicted visual properties of an upcoming reach during planning. It is still an open question whether such coding might is also present in human PPC and, moreover, whether such visual representations may also exist in other regions of the brain, for instance in premotor cortex.

The latter seems a valid question as dorsal premotor cortex (PMd) has been shown to closely work together with PPC in several aspects of action planning, execution and monitoring (Desmurget and Sirigu, 2009; Hoshi and Tanji, 2000; Westendorff et al., 2010; Lindner et al., 2010) Moreover, as demonstrated already, PMd prospectively encodes not only hand-target vectors in visual coordinates (Pesaran et al., 2006; Ochiai et al., 2002) but also any initial direction of movement to circumvent obstacles while reaching (Pearce and Moran, 2012). These findings yield some resemblance to the described functions of the posterior parietal cortex in representing movements withing a sensory reference frame (see e.g. Andersen and Buneo, 2002). On the other hand, several lines of research suggest, that the frontal areas represent movements more in body, than in sensory space, as opposed to posterior parietal cortex (Beurze et al., 2010). The available data draw therefore a rather complex picture of planning processes in PPC and PMd and do not make it clear how are the sensory properties incorporated into movement plans by these areas.

Here we attempted to investigate the relationship between the visual consequences of an action and their underlying motor plan representations in PPC and PMd. For this purpose we designed a virtual reach task, in which we systematically manipulated the interrelation between movements and their visual outcome by altering the visual movement feedback while keeping the motor demands constant. Using this approach we asked whether planning activity in human PPC and PMd does indeed contain a representation of the desired visual consequences of upcoming goal-directed movements, independent of the motor programs producing them.

## MATERIALS AND METHODS

### Subjects

14 healthy subjects (8 females) participated in the study. All of them had normal or corrected to normal vision. All except one subject were right-handed. All participants gave written informed consent prior to participation in the study. The experimental procedures were carried out in accordance with the declaration of Helsinki, and the study was approved by the ethics committee of the University Hospital and the Faculty of Medicine at the University of Tuebingen. The participants and were reimbursed for their participation. Two of the participants were excluded from the final sample (see “Behavioral performance analysis” for details).

### General task design

To study planning-related brain activity we conducted a functional magnetic resonance imaging (fMRI) study, in which human subjects performed an action planning experiment (Figure 1). During this experiment subjects needed to plan and execute “virtual reaches” by moving a button-controlled cursor on a response-grid. In half of the trials, subjects carried out a delayed response task (Rosenbaum, 1980): they were instructed to remember a target location presented during the initial cue epoch and plan a movement towards it. Then, after an intervening delay epoch during which the target was no longer present, they had to execute the pre-planned movement during a movement epoch. In these trials, it was necessary to plan a movement prior to the movement epoch, hence we named this task “*pre-planned movement task*” (**PPM**). In the other half of the trials, subjects were told to ignore the initial cue and instead to wait until the movement epoch of that trial. Then they had to move the cursor to a new, visually instructed target location, randomly placed on the response grid. The latter task was named “*direct movement task*” (**DM**) and differed from the PPM in that both movement planning and execution took place directly during the movement epoch. Contrasting both types of trials allowed us to access brain processes related to movement planning. First, comparing delay-related brain activity in PPM vs. DM should allow one to isolate activity due to movement pre-planning in PPM (Rosenbaum, 1980; Lindner et al., 2010). Second, contrasting the estimates of brain activity during the movement epoch for DM vs. PPM should exhibit activity related to initial, fast planning processes that still need to be accomplished in DM but that are already completed in PPM (c.f. Ames et al., 2014).

**Figure 1.**
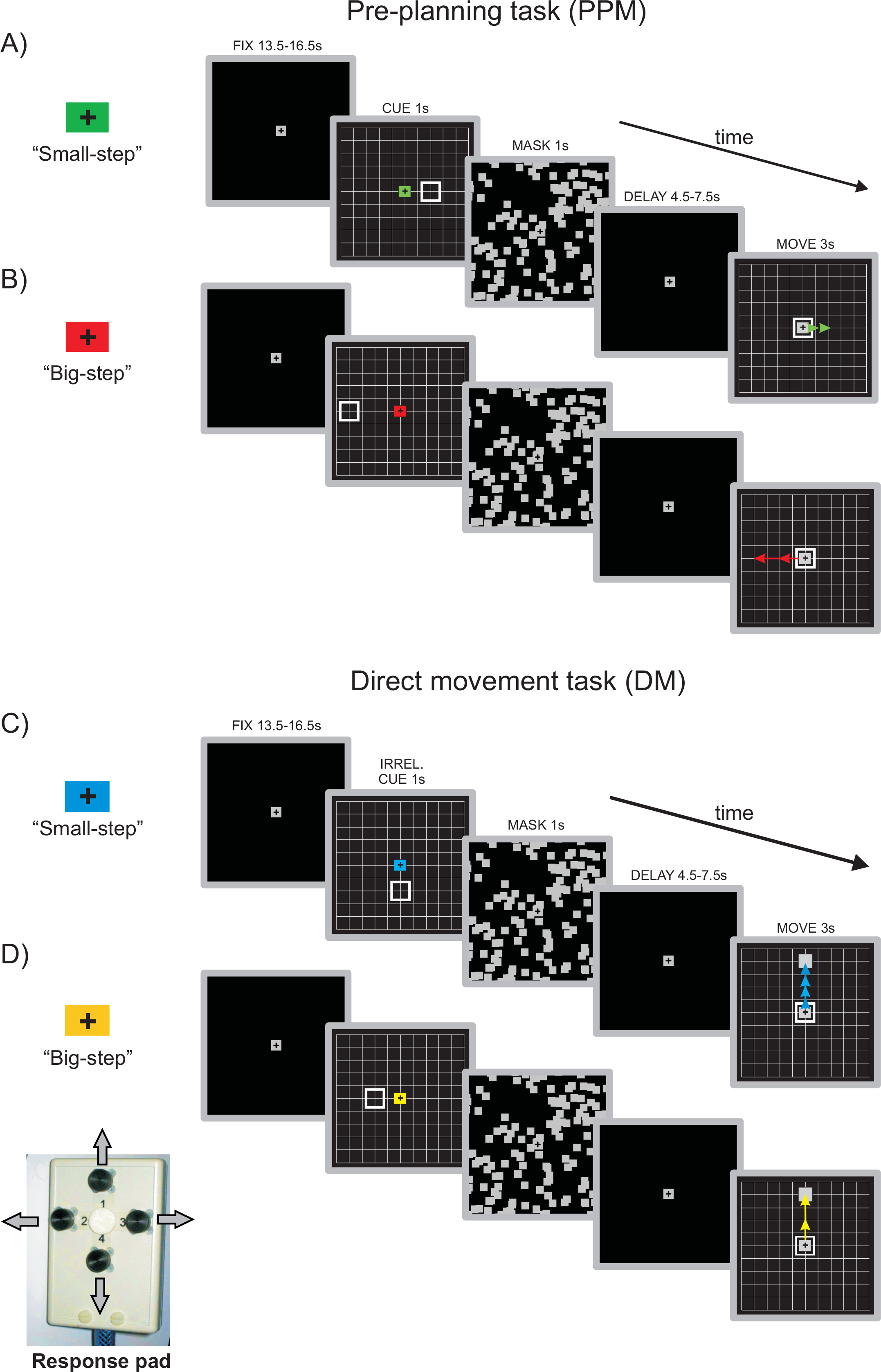
Experimental Tasks. Schematic trial timelines showing each combination of task (DM, PPM) and movement gain (“small step”, “big step”). Each trial started with a baseline-fixation period (FIX). Next, a grid-like movement space was presented for 1s, containing a target location in the periphery and a color cue in the center indicating subjects the specific task and gain context in a given trial (CUE). Next the screen was masked for 1s (MASK) and afterwards only the fixation cross remained visible, indicating the delay period (DELAY). When the grid appeared again (MOVE), subjects were required to perform a movement to the remembered pre-cued target location in PPM trials (**A & B**) or to the newly presented grey target in DM (**C & D**). Arrows (not visible to the subjects) illustrate the way that the virtual end effector moved after each single press of the button given the specific gain context, performing either a “small step” ( **A** & **C**) or a “big step” movement (**B & D**). After a time limit of 3s the movement screen was masked again and the subsequent trial began.

To further address whether any of the planning activity revealed would reflect the desired visual outcome independent from the motor components of a movement, we additionally manipulated the visual movement gain of the cursor in both tasks. This meant that after each single button press, the cursor could perform either a “small step” (i.e. jump to the next intersection of the response grid) or a “big step” movement (i.e. jump to the next, and then to the second-next intersection). Targets were positioned in such a way that they required sequences of 1, 2 or 4 button presses to be reached, in each of the “gain” conditions. By changing the movement gain ("big" or "small" step) for each given movement sequence length, we could keep constant a sequence’s motor demands while at the same time vary its visual consequences (the visual distance of the movement). This was meant to allow us for capturing planning activity that would specifically reflect the amount of upcoming visual motion. Since the number of trials with each sequence length was balanced, the only difference between the "big" and "small step” conditions in each task was the amount of visual motion the sequences produced.

Information about which movement gain was actually applied in a given trial was shown to participants during the cue epoch of the trial (see Figure 1) and they needed to incorporate this information into their motor plan in order to perform accurately within the time limit of the movement epoch. For every participant, color cues indicating conditions were the same.

Using our approach we expected to reveal a representation of the upcoming motor sequence length during movement planning in PPM, a representation that has already been described previously for both PPC and PMd (Lindner et al., 2010). Areas that would exhibit such a prospective representation of the motor sequence were considered in a subsequent region of interest (ROI) analysis (see below), in order to reveal whether their planning activity contains additional information about the visual consequences of an upcoming movement. In other words, the planning activity should represent the “visual way to the goal” in addition to the motor sequence. Previous fMRI findings revealed that the amplitude of the blood oxygenation level dependent (BOLD) signal correlates positively with the amount of (anticipated) visual motion (Lindner et al., 2006). Therefore we hypothesized that if the visual consequences of a planned movement are indeed defined in PPC or PMd, the brain activity in these areas should reflect these visual aspects of the movement. Specifically, the "big-step" motor sequences should on average produce stronger planning-related BOLD signals as compared to the "small-step" sequences, due to an overall larger amount of expected visual motion.

### Stimulus presentation

Stimuli were presented using Cogent Graphics Toolbox (Laboratory of Neurobiology at the Wellcome Department of Imaging Neuroscience London, UK) running on a Windows^TM^ based PC and delivered to the subject using a LCD projector (1024×768 pixels, 60Hz refresh rate), a translucent screen and a set of mirrors attached to the head coil of the MRI scanner.

Each trial started with a baseline epoch (13500, 15000 or 16500ms), which required the subject to fixate their gaze upon the centrally positioned fixation cross (1.1 deg visual angle). The fixation cross remained visible for the whole time course of a trial and subjects were instructed to fixate it at all times. This should help us to avoid, potentially confounding, eye-movement related brain activity. After the baseline epoch ended, the cue screen was presented for a fixed time of 1000 ms. The cue screen consisted of the movement space grid (9×9 squares, see Figure 1; angular size of each square was approx. 1.7 deg) and an empty square representing target location (approx. 1.7 deg). The fixation cross was replaced by a color cue, that indicated both the movement gain (“big-step” vs. “small-step” movement) and the task (PPM or DM). Subsequently the scene was masked for 1000ms to prevent afterimages of the visual targets and then the delay epoch began. The delay epoch was of variable length (4500, 6000 and 7500ms). The delay times chosen were much shorter than those used in most previous studies (e.g. compare Lindner et al., 2010). We did this to reduce subjects ability to use retrospective mnemonic strategies and from engaging in task-unrelated cognitive activities (such as any sort of mind-wandering) during the delay epoch. Subsequently, the response grid was presented, and subjects were supposed to execute the movement to the remembered target location (in case of PPM) or to a filled square (1 deg), indicating the actual target in DM. Specifically, subjects used a response pad (see Figure 1) held in their right hand in order to move a button-controlled square cursor to the designated target area. The cursor moved in the direction that corresponded to the button pressed (either left, right, up or down), skipping between intersections along the vertical or horizontal lines of the grid, respectively. Depending on the movement gain, a single press of the button could either lead to a “small-step” movement of the cursor (so that the cursor moved from one intersection on the grid to the next) or a “big-step” (in this case the cursor jumped twice in the same direction, with a 100ms time delay between successive cursor steps). Time for completing the motor response was limited to 3000ms. After the time limit was reached, the screen was masked again, and the next trial began after an inter-trial interval of 2000ms.

Subjects practiced the task before scanning. During scanning subjects performed 36 trials for each of our four conditions in total. These trials were acquired during three experimental blocks, each consisting of 48 trials. The conditions were randomized within a block.

### Oculomotor behavioral control

Eye movements were monitored at 50Hz sampling rate with an infrared operated, MR-compatible eye tracking camera (SensoMotoric Instruments) and the ViewPoint software (Arrington Research). All eye movement analyses were performed off-line using custom routines written in Matlab (MathWorks). In brief, eye position samples were filtered using a second-order 10 Hz digital low-pass filter. Saccades were detected using an absolute velocity threshold of 20 degrees per second.

Since our experiment required subjects to maintain fixation on the central fixation point, we excluded 2 of our 14 participants who did not comply to this instruction and frequently performed large saccades (amplitude > 3 degrees visual angle away from the fixation point in more than 20% of the trials). In both excluded subjects, this behavior was equal to gaze shifts towards the target location in the cue epoch, or towards the moving cursor during the movement epoch of a trial, or towards both.

### Manual performance analysis

Manual performance was assessed in terms of hit rate, movement durations and reaction times. Those trials were classified as hits, in which the cursor was positioned over the correct target location at the end of the movement epoch. Movement duration captured the time from the first button press until the cursor reached its final position. Reaction time was defined as the interval between the onset of the movement epoch and the time at which the first button in a sequence was pressed.

All behavioral data were analyzed statistically using 2×2 repeated measures ANOVA with factors “task” and “movement gain”.

### fMRI acquisition and SPM analysis

MRI images were acquired on a 3T Siemens TRIO scanner using a twelve-channel head coil (Siemens, Ellwangen, Germany). For each subject, we obtained a T1-weighted magnetization-prepared rapid-acquisition gradient echo (MPRAGE) anatomical scan of the whole brain (176 slices, slice thickness: 1 mm, gap: 0 mm, in-plane voxel size: 1 × 1 mm, repetition time: 2300 ms, echo time: 2.92 ms, field of view: 256 ×256, resolution: 256 × 256) as well as T2*-weighted gradient-echo planar imaging scans (EPI): slice thickness: 3.2 mm + 0.8 mm gap; in-plane voxel size: 3 × 3 mm; repetition time: 2000 ms; echo time: 30 ms; flip angle: 90°; field of view: 192 × 192 mm; resolution: 64 × 64 voxels; 32 axial slices. Overall, we obtained 2100 EPIs per subject, which were collected during the three consecutive runs of about 20 min length each. A single EPI volume completely covered the cerebral cortex as well as most subcortical structures. Only the most inferior aspects of the cerebellum were not covered in several of our subjects.

Functional data were analyzed using SPM8 (Wellcome Department of Cognitive Neurology, London, UK). In every subject, functional images were spatially aligned to the first volume in a series, and then coregistered to the T1 image. After that, a non-linear normalization of the structural image to a T1 template in MNI space was performed. Parameters obtained with this normalization were then applied to all functional images. In the last step of data preprocessing, we smoothened all the functional images with a Gaussian filter of 8 mm × 8 mm × 8 mm FWHM.

In subject-specific fMRI analyses we specified two general linear models for each individual. The first model included all the four conditions (“task” × “movement gain”) and for each condition we modeled trial epochs (cue+mask, delay, response) as separate regressors. Cue was modeled as a single regressor, regardless of the condition. Sequence length was modeled as a linear parametric modulator, thus capturing any relative difference in BOLD-signal amplitudes related to the number of button presses required to reach targets at different distances. This parametric modulator was included separately for all conditions and for both delay and movement epoch. Head motion parameters were included in the model as six independent regressors (x, y, z translation and x, y, z rotation). Inter-trial intervals, as well as fixation epochs weren’t modeled explicitly and thus served as an implicit baseline.

The second GLM was constructed in order to obtain reliable cue-related betas for each of the experimental conditions (see “results”). This would allow us to scrutinize early part of planning processes in the PPM. To this end we modeled cue epoch in the same way as other epoch regressors, namely defining “task” and “movement gain” separately. The other regressors we modeled as described above.

### ROI analysis

For each subject we identified a set of regions that contributed to the prospective planning of motor sequences (Figure 2; Compare: Table 1). We decided for this region-of-interest (ROI) approach in order to avoid inter-individual variation in functional anatomy and focus on the planning-related activity in the relevant areas. Towards this end we first calculated the statistical parametric map capturing areas that show a parametric modulation of their BOLD activity by the planned number of button presses (motor sequence length) during the delay epoch of “pre-planning” trials in each individual. Based on coordinates of movement sequence planning regions that were described by Lindner et al. (2010), we selected our ROIs by looking for areas showing a statistically significant linear increase in BOLD intensity during the delay phase of PPM trials within a search radius of 20mm around the respective coordinates (p<0.05, FWE-corrected for multiple comparisons within the search volume). These areas were: left and right superior parietal lobule (SPL), left and right dorsal premotor cortex (PMd) and anterior intraparietal sulcus (aIPS). Since for the left aIPS we were only able to identify these areas in 9 out of 12 subjects, we applied a more liberal threshold (p<0.001 uncorrected) in the remaining 3 subjects to provide a more representative sample. In addition to these planning ROIs we included several control ROIs: (i) The hand representation in the left and right primary motor cortex (M1) was identified based on anatomical criteria (Yousry et al., 1997) to control for motor response-related activity and for effector preparation (ii) dorsolateral prefrontal cortex (DLPFC) was mapped according to the same criteria as described for our planning ROIS. Like for aIPS, a more liberal threshold (p<0.001 uncorrected) was applied in three subjects. In only one subject we were not able to reliably localize DLPFC using the latter criterion. Data from that subject’s DLPFC were therefore extracted using group-based coordinates. (iii) Finally, we additionally included area V1 as a control ROI in order to capture activity reflecting visual input stemming from the target cue or the cursor movement. We we are not aware of any findings showing its specific engagement in reach planning.

**Figure 2.**
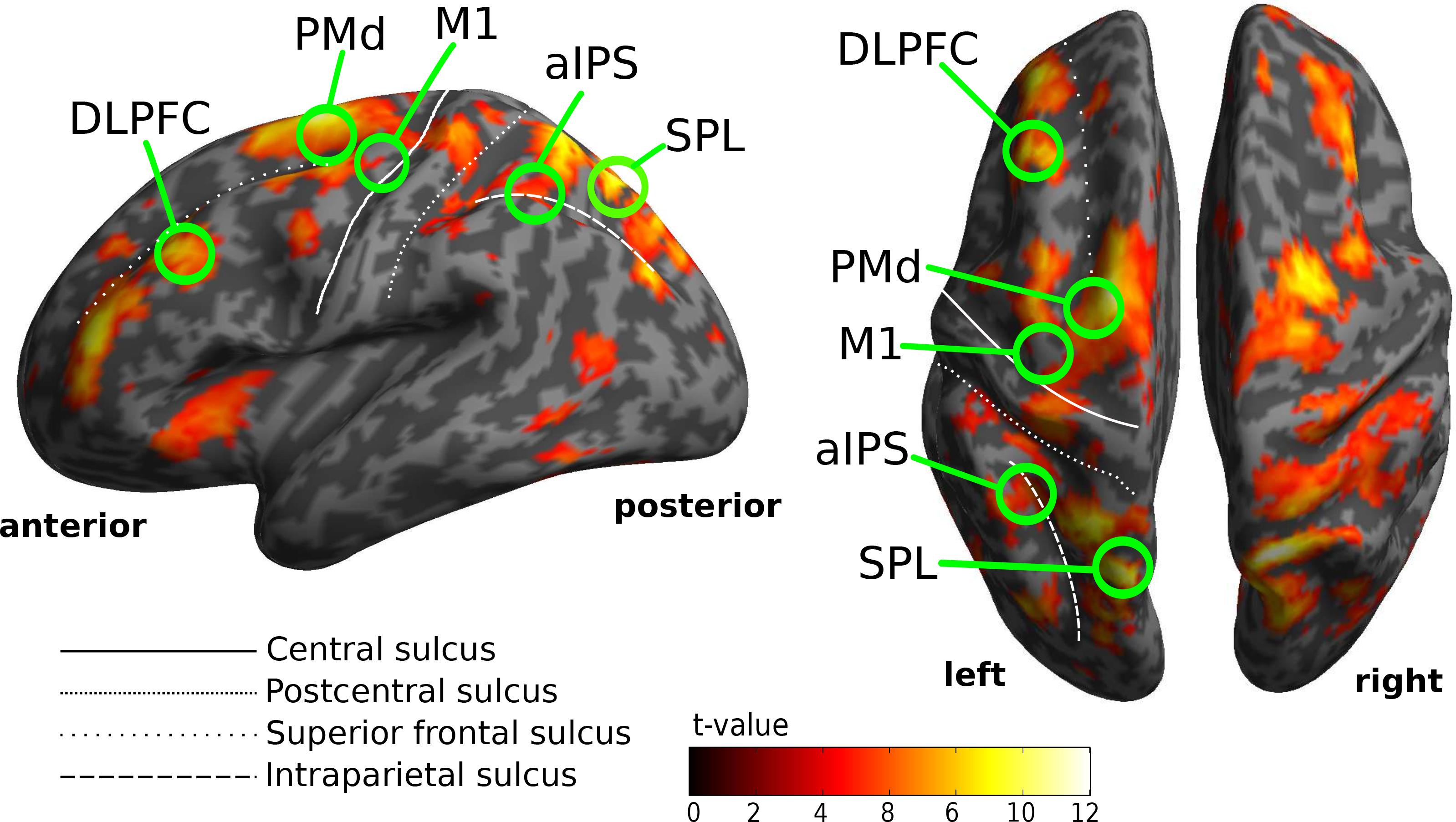
Areas of planning-related fMRI activity representing motor sequence length in an exemplary subject. The statistical parametric map is thresholded at p<0.05, FWE corrected for multiple comparisons (see “MATERIALS AND METHODS” for a detailed description of the ROI selection criteria).

**Table 1.**
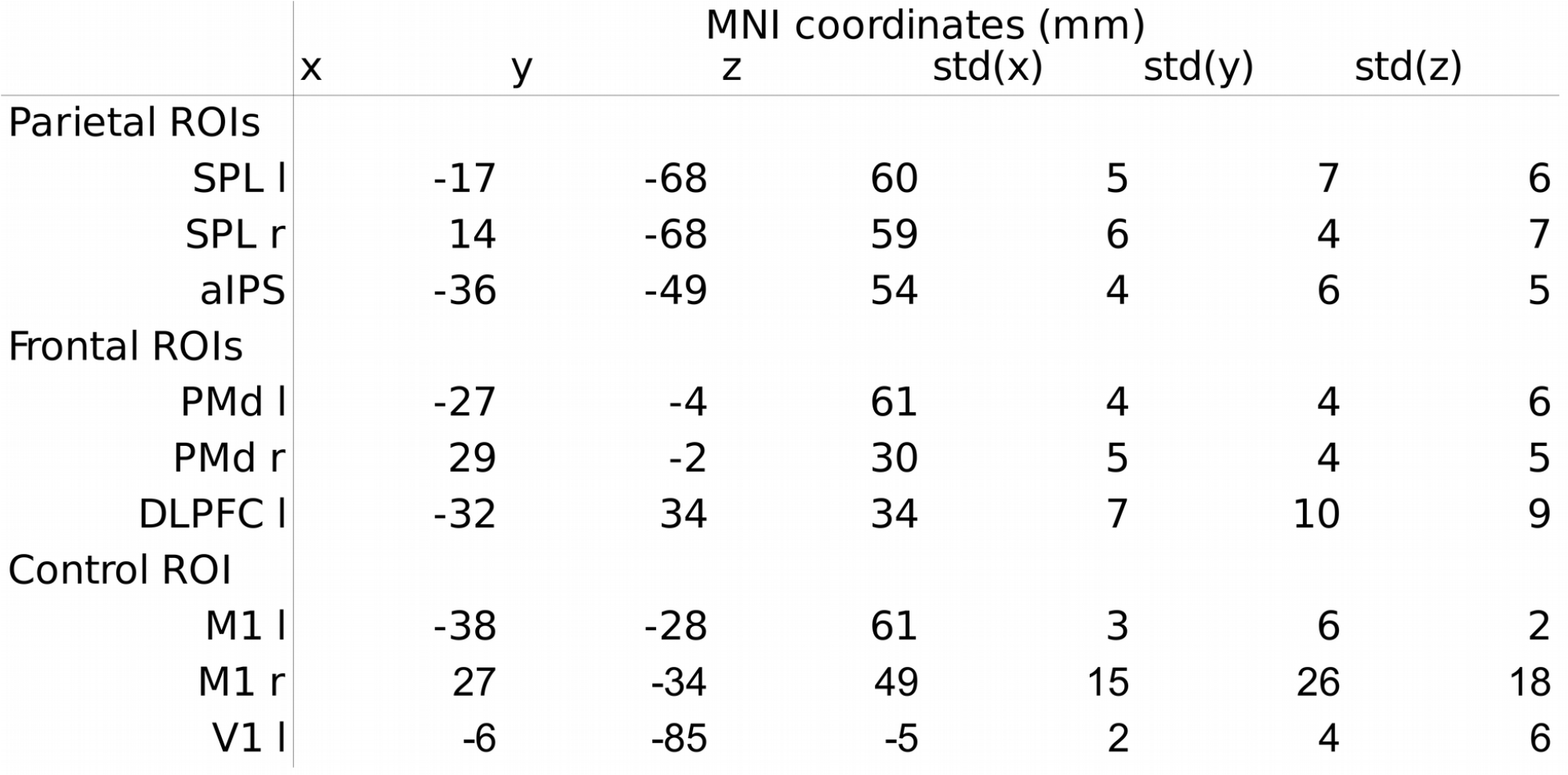
Average locations of ROIs.

As our functional ROI definition did not differentiate between “movement gain” conditions (i.e. maps were calculated for both “small-step” and “big-step” movements taken together), the ROI selection was not biased in favor of our hypothesis (i.e. stronger planning activity in “big-step” conditions). In the next step, for each of our ROIs and in each of our subjects, we extracted the normalized mean beta weights of our main GLM regressors from a 3mm radius sphere created around the ROIs center coordinate, each single ROI consisting of 7 voxels in total. The extracted betas of the first GLM were analyzed separately for the delay and the movement epoch using a 2x2 repeated measures ANOVA with the factors “task” and “movement gain”. The GLM was analyzed using the added additional factor (“epoch”) in a 2×2×2 repeated measures ANOVA.

## RESULTS

### Behavioral performance

We controlled several behavioral variables relevant for the interpretation of our fMRI data (Figure 3). Specifically, to demonstrate that subjects prepared their movement plans prior to the movement epoch in PPM trials, we analyzed subjects’ reaction times (see “Experimental procedures” for details). Reaction times in PPM trials were contrasted to those revealed in DM trials, as in the latter trials planning could take place only in the movement epoch (i.e. after the target had been presented) allowing us to estimate the reaction time benefit through pre-planning (Rosenbaum 1980). As expected, manual reaction times were on average significantly shorter in pre-planned movement trials (PPM) than in direct movement trials (DM) (2×2 repeated measures ANOVA, main effect “Task” (PPM vs. DM): p<0.001; main effect “Movement gain” (“small-step” vs. “big-step”): p>0.05, n.s.; interaction: p>0.05, n.s.) (Figure 3A).

**Figure 3.**
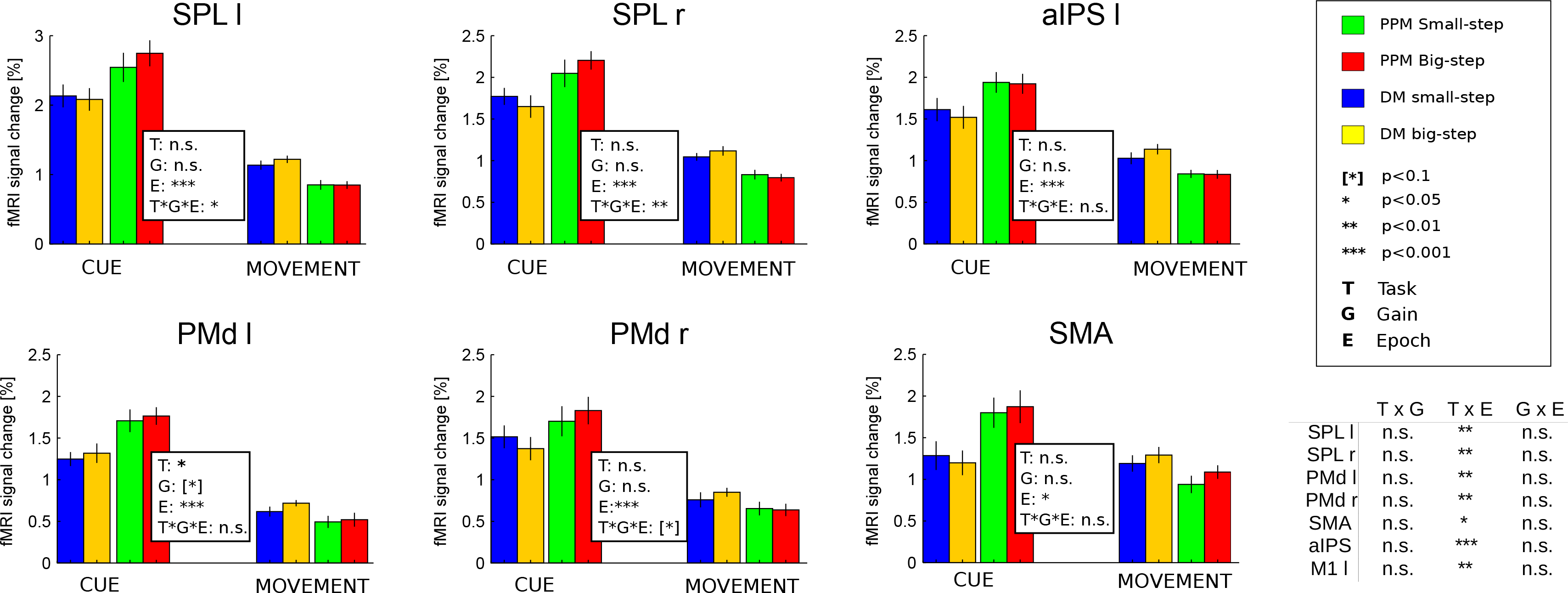
Behavioral Performance. Average manual and oculomotor performance in PPM and DM trials. Error bars denote standard deviations in A)-C) and standard errors in D). **A** Manual reaction times in PPM trials were significantly shorter than in DM trials, indicating that planning resulted in a reaction time benefit in PPM. **B** Hit rates did not differ significantly between PPM and DM, nor between movement gains, implying balanced movement difficulty. **C** Movement durations were on average significantly longer in “big-step” trials than in “small-step” trials, regardless of task. This difference was explained by the longer movement animation of the visual end-effector in “big-step” trials (see “Experimental Procedures”) **D** The frequency of fixational saccades during delay and movement epochs did not differ between PPM and DM and also did not vary with gain context. In the cue epoch there were significant effects of “movement gain” and “gain”x”task” interaction.

Hit rates were constant across both tasks and movement types (2×2 repeated measures ANOVA, main effect “Task”: p>0.05, n.s.; main effect “Movement gain”: p>0.05, n.s.; interaction: p>0.05, n.s.), indicating that movement difficulty across conditions was balanced (Figure 3B).

Average movement durations showed an expected effect for the factor “movement gain” (i.e. “big-step” sequences produced significantly longer durations because of the way the cursor movement was animated (see “Experimental Design” for details), but no “task” and interaction effects were present (2×2 repeated measures ANOVA, main effect “Movement gain”: p<0.001; main effect “Task”: p>0.05, n.s.; interaction: p>0.0.5, n.s) (Figure 3C).

An eye movement data analysis yielded no significant difference between both tasks and movement types with respect to the number of fixational saccades during delay and movement epochs (2×2 repeated measures ANOVA, main effect “Movement gain”: p>0.05, n.s.; main effect “Task”: p>0.05, n.s.; interaction: p>0.05, n.s) (Figure 3D). This ensured that fMRI activity in these epochs was not differentially influenced by varying oculomotor behavior across conditions. The saccade rates were, however, significantly different across conditions in the cue phase, as an ANOVA revealed a significant main effect of “Movement gain” (F=5.056, df=11, p=0.046) and a “Task”×”Movement gain” interaction (F=11.694, df=11, p=0.006). As we will discuss later in the text, these effects cannot explain the reported fMRI results.

### Planning activity encodes visual properties of upcoming movement

For studying planning-related brain activity we decided for region of interest (ROI)-based approach to focus on the areas that were previously demonstrated to contain prospective representations of motor sequences (Lindner et al. 2010) and these motor representations we assumed likely to be modulated by expected visual properties of actions. First, we performed a whole-brain analysis in single subjects to define for each subject the brain areas that exhibited significant modulation of planning activity by motor sequence length during the delay epoch of PPM trials (see “Materials and methods” for details; also compare: Lindner et al. 2010). The following regions exhibited such modulation of planning activity in all subjects: superior parietal lobule (SPL, bilateral), dorsal premotor cortex (PMd, bilateral) and anterior intraparietal sulcus (aIPS, left) (Figure 2). On the basis of previous research (Lindner et al., 2010) we assumed that such activation pattern is characteristic for areas contributing to the prospective planning of goal-directed motor sequences and that – in a second step – we could test whether activity in these ROIs is modulated by movement gain. In addition, we included dorsolateral prefrontal cortex (DLPFC, left), the hand area of left and right primary motor cortex (M1) and area V1 as additional control ROIs. It is worth to emphasise, that our functional ROI selection criterion was independent to the tested hypothesis and thus allowed us to avoid circularity in subsequent analyses.

ANOVAs performed on the activity estimates (i.e. the normalized beta weights) extracted from these ROIs for the movement phase revealed a significantly stronger BOLD signal in DM than in PPM in several areas, namely left and right SPL, left and right PMd, and M1 (Figure 4). We consider these task-related changes an indicator for planning processes in DM: any pre-planning during the delay would strongly reduce the cognitive load needed to plan and execute actions in the movement phase of PPM. On the other hand, planning was still needed during the movement phase in DM (Ames et al., 2014), thus elevating related BOLD signal amplitudes in DM as compared to PPM.

**Figure 4.**
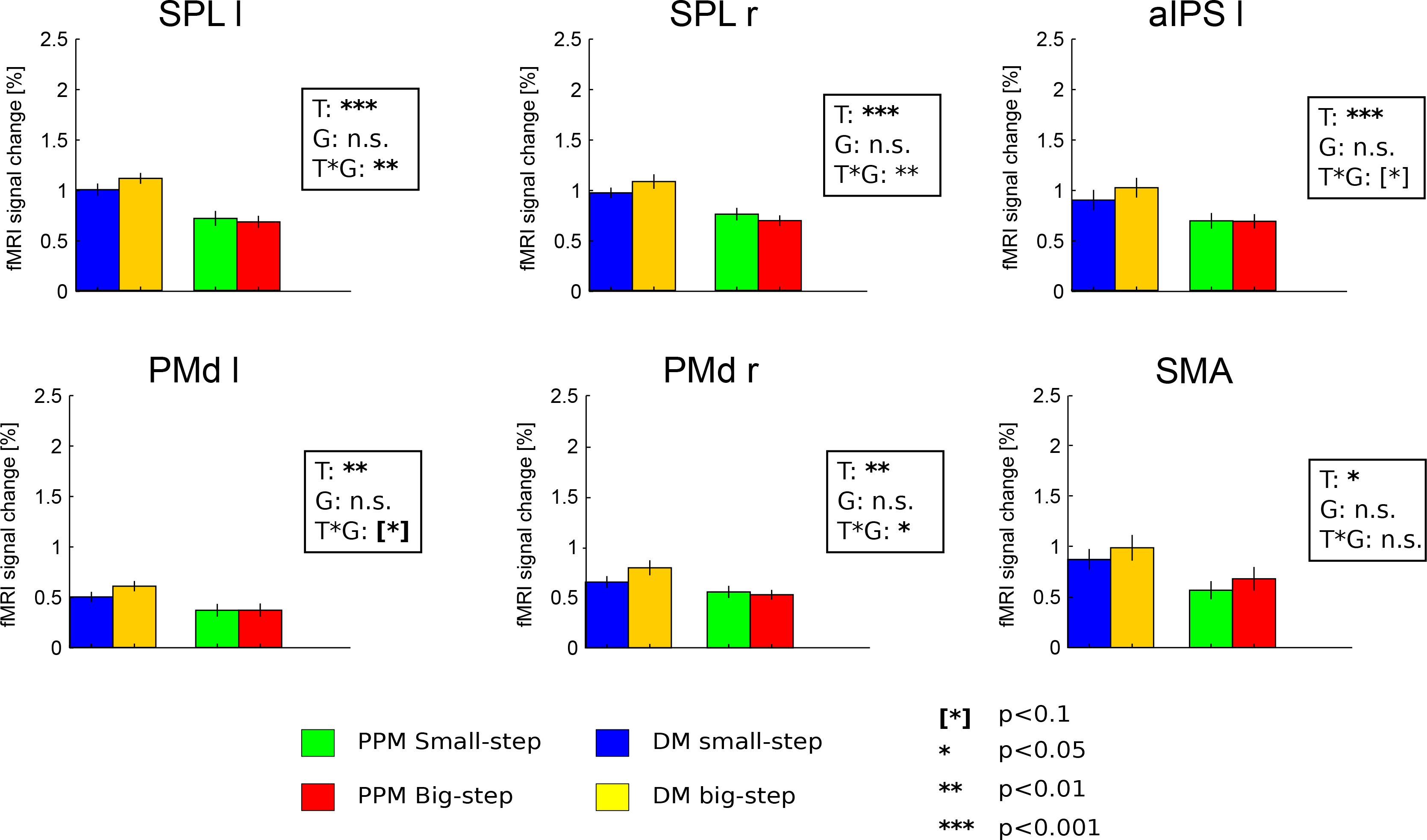
BOLD activity in all ROIs during the movement epoch. Signal increases in DM with respect to PPM reflect planning processes (compare main text). Signal differences between “big-step” and “small-step” trials in left and right SPL and in right PMd refer to an influence of (planned) visual movement distance. All values represent averages calculated across subjects’ mean activity +/−SEM.

Most importantly, in both medial PPC and PMd putative planning activity in the movement phase of DM was additionally modulated by movement gain: the “big-step” motor sequences elicited on average significantly higher BOLD signal amplitudes than did the “small-step” sequences in left and right SPL and right PMd, as indicated by the significant interaction of the factors “Task” and “Movement gain” (Figure 4). This indicates that the visual aspects of upcoming movements were represented in these regions. It is noteworthy that there was also a nearly significant trend for this effect in left PMd (p=0.088), implying a bilateral representation of the visual movement consequences in that area as well. A similar trend was also observed in left aIPS (p=0.09).

No such movement gain-related pattern was present in the movement epoch of PPM in these (and all other) ROIs, indicating that it is not the visual motion *per se* that would explain the signal differences between “big step” and “small step” movements (Figure 4).

We neither did observe a gain-related modulation of brain activity in the movement epoch of DM in primary motor cortex nor in dorsolateral prefrontal (Figure S2). Area M1 is primarily engaged in preparation and execution of motor programs (see eg. Hocherman and Wise, 1991), and, at least to our knowledge, there is no evidence that it could process any visual information about the upcoming action. The dorsolateral prefrontal cortex, in turn, has been demonstrated as being engaged in retrospective mnemonic processes rather than in prospective planning (e.g. compare Lindner et al. 2010). Therefore the lack of signal modulation due to movement gain in these particular ROIs additionally supports our main hypothesis (see DISCUSSION).

It is also worth to note that both in PMd (in PPM and DM trials) and PPC (in DM trials) visual modulation of planning activity was easier to observe (see p values) on the ipsilateral but not the contralateral side (with respect to the effector). This suggests that ipsilateral representations of movement may organize information more in terms of the abstract (visual) motor plan, whereas the contralateral representations might process information in a way that more directly refers to effector’s motor action. While we cannot reliably test this hypothesis on the basis of the current dataset, it seems at least to be supported by findings of Krasovsky et al. (2014) who also reported that representations of sensory action-outcomes are rather ipsi-, than contralaterally organized.

Comparisons of activity estimates for the delay epoch did neither reveal any significant gain-related differences in any of our planning ROIs nor any differences between movement tasks, i.e. PPM vs. DM (see Figure S1). The lack of a difference between PPM and DM during the delay, which contrasts previous studies (e.g. Lindner et al. 2010), could result from the comparatively short delay epochs in our study. This suggests that such short delays apparently do not allow a full separation between the cue- and delay-related BOLD-signals and can be also susceptible to any instruction-independent default planning in (Snyder et al., 2006) in response to the irrelevant target cues that were presented during the cue epoch of DM trials. Irrespective of these limitations of our design with short delays, one may still suspect that the difference in movement gain could still be reflected by sustained BOLD-signals in the delay phase of PPM. In our view, our inability to observe such effect demonstrates one potential weakness of using the classical delayed response paradigm in fMRI research, namely its limited capacity to capture brain responses related to rapid, early planning processes. This speculation is supported by aforementioned findings of Kuang et al. (2015) who demonstrated, that the relative amount of visual planning neurons is significantly higher early during the planning stage of an action and is becoming less pronounced later during the delay, in the sustained neural response. We therefore find our approach of focusing at DM activity, a valid alternative to study rapid planning processes.

Certain delay-related differences between tasks were present in control ROIs: Interestingly, activity in the left motor cortex was significantly stronger in the PPM than in DM (p=0.0027), apparently reflecting unspecific effector preparation processes. This is confirmed by the lack of such modulation on the ipsilateral side (see Supplementary Figure S2). Likewise, in DLPFC there was a significant influence of task, namely a stronger activity in PPM as compared to DM too. This area might be engaged in mnemonic aspects of motor planning (e.g. a retrospective representation of the movement target), as was suggested by previous findings of Lindner and colleagues (2010).

The lack of a gain effect in activity of planning ROIs during the delay epoch in PPM trials prompted us to look more closely at early planning activity in these trials. This is because early planning processes might be already reflected in the integrated BOLD-signal during the cue-epoch. Therefore we wanted to contrast such early planning in PPM during the cue epoch with early planning in DM during the movement epoch. For this purpose, we estimated an alternative GLM in which we now also focused on gain-related changes during the cue epoch (as compared to the response epoch). We ran a three-way repeated measures 2×2×2 ANOVA with the factors “Task” (PPM and DM), “Movement Gain” (“small-” and “big-step”) and “Epoch” (“Cue” and “Movement”). We assumed that the visual effect (“big-step” > “small-step”) should be visible in the cue epoch of PPM and in the movement epoch in DM, as would be confirmed by a three-way interaction of the three factors. Indeed, this analysis uncovered that the early planning response in left and right SPL do show the expected effect of visual movement properties as dependent on task and trial epoch (Left SPL: F=5.469, df=11, p=0.0393; right SPL: F=10.946, df=11, p=0.0070). In right PMd we revealed a clear trend for the same effect (F=4.476, df=11, p=0.0580). The gain effect was however absent in left PMd (F=0.249, df= 11, p=0.6273). This finding shows that the visual aspects of movement are indeed present in the early planning activity.

## DISCUSSION

### Prospective representation of visual movement consequences

Our experiment demonstrates an alternative approach to studying planning-related fMRI-activity of the human brain. Instead of only focusing on delay epoch between instructive cue and action initiation (e.g. compare Lindner, 2010), we chose to compare movement sequences that had been already pre-planned (PPM) to those that required fast planning directly before execution (DM) (compare Ames et al., 2014).

Using this approach, we were able to observe an additional modulation of DM activity by the visual consequences of movement in the same ROIs. Activity was the stronger the more visual motion the same movement sequences produced due to the gain manipulation. This effect was apparent in areas previously demonstrated to contain prospective representations of action plans (PPC and PMd; see: Lindner et al. 2010). Yet, it was absent both in primary motor cortex and in DLPFC. The latter have previously been demonstrated to maintain a retrospective memory of visual movement targets (Lindner et al., 2010). The modulation of planning activity by movement gain was present also in early cue-related brain responses but not during the delay period of PPM trials. This suggests that sensory representation are inherent to earliest stages of movement planning, where processes like target localization and movement path definition crucially depend on vision. Once this early plan is defined, relevant motor programs are constructed and remain maintained in memory.

Our results are therefore consistent with the idea that motor planning activity in PPC and PMd initially represents the visual consequences of an upcoming movement while the required motor programs needed to realize such visual action plans arise at later stages of sensorimotor processing and it seems likely that only these motor programs are maintained in memory until being ultimately put into action.

### Alternative paradigms for dissociating vision and manual action

Apart from the gain manipulation that was applied in our study, other experimental paradigms have been used to alter the interrelation between hand movements and visual information. These paradigms could potentially provide us with additional clues about how the visual consequences of manual actions are embedded in an action plan. One such paradigm is the so-called anti-reach task in which subjects need to perform reaches towards a location opposite to a pre-cued visual target location (Crammond, Kalaska, 1994; Westendorff et al. 2010). While this task clearly allows distinguishing activity related to the direction of a visual target vs. activity related to the direction of movement, it cannot discern whether any movement-related activity would refer to the visual or to the bodily direction of movement as both are identical. Another class of paradigms that seems related engages inverting prisms (Helmholtz, 1909; Clower 1996). The use of prisms can clearly help to dissociate bodily motion from its visual consequences (e.g. through inverting prisms). Yet, when monitoring brain activity during such paradigms particular care has to be taken to disentangle whether activity truly reflects the visual consequences of movement rather than any visual stimulus itself (or the memory thereof), as visual movement- and target-direction are identical. So far there is only one electrophysiological study on action planning in monkeys that has combined both paradigms and that therefore could account for the aforementioned limitations (Kuang et al., 2015; for details see “Introduction”). In our own human fMRI experiment the visual movement consequences and the location of the visual goal were also tightly coupled, but our specific experimental findings still allowed us teasing apart these factors as will be discussed in the following paragraph.

### Potential limitations of interpretation

Before answering what action components determined the gain-related modulation of the BOLD signal in the early planning activity, some potential confounding factors need to be considered.

In our eye movement analysis we revealed a significant influence of experimental condition on saccadic frequency but during the cue phase, only. Here, saccades were most frequent in the “small-step” DM condition. Therefore, when assuming that saccade rates are positively correlated with the amplitude of the BOLD response (see eg. Kimmig et al., 2001) this saccade effect can hardly account for the pattern of gain-dependent planning activity in PPC and PMd during the cue phase, namely the change in PPM-related activity (compare Figure 3D and Figure 5). Moreover, as there was no difference in saccade rates in the movement phase, saccadic eye movements also cannot explain the gain-related modulation of planning activity in DM during this task epoch.

**Figure 5.**
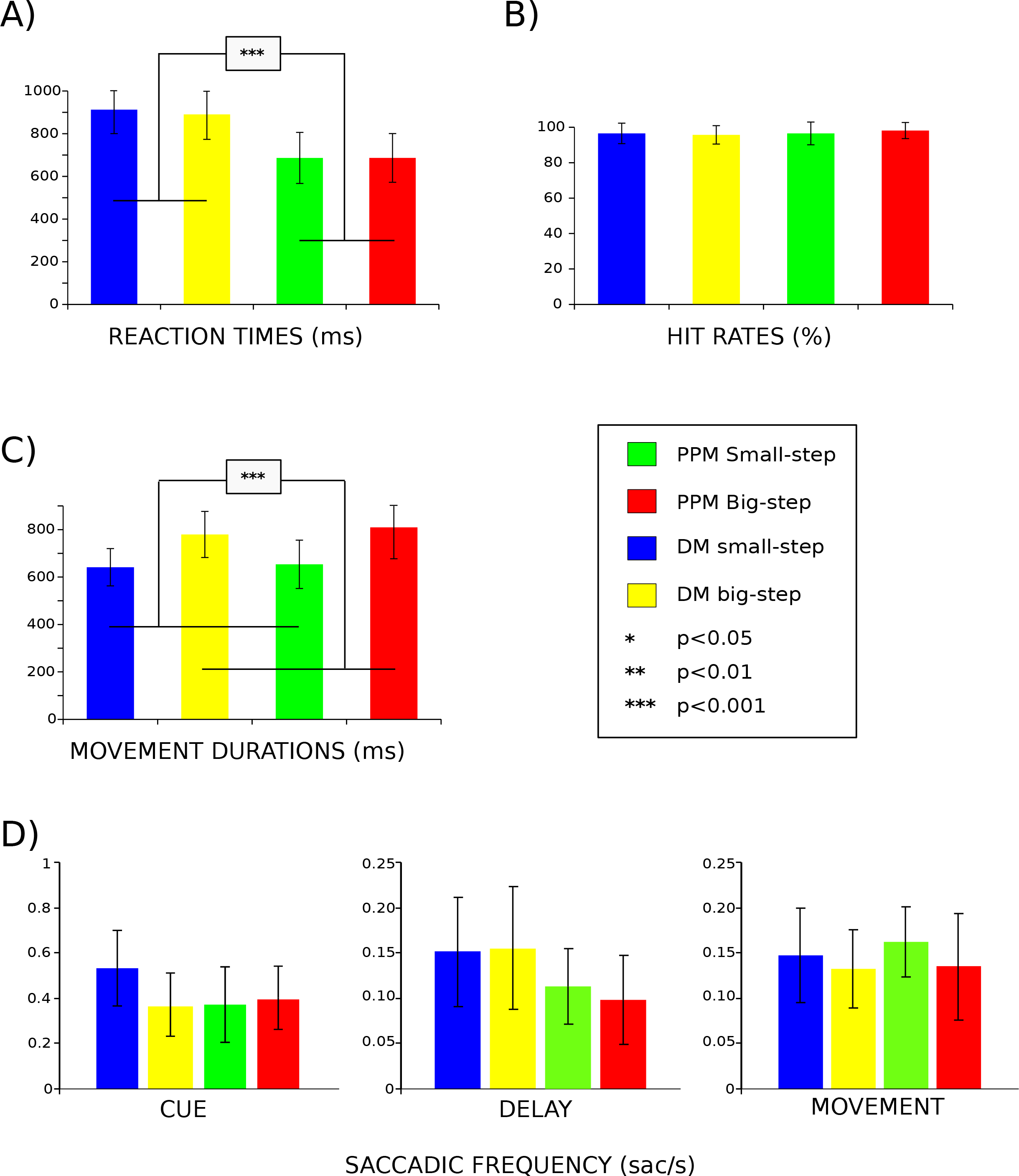
BOLD activity at early planning stages. Cue epoch of PPM and movement epoch of DM are grouped together as “planning” conditions. The Task(T) × Gain(G) × Epoch(E) interaction demonstrates presence of visual consequences representation in the left and right posterior parietal cortex (SPL). A significance-nearing trend is present also in the right PMd. The additional table contains interaction effects for all the depicted ROIs. Task × Epoch interaction demonstrates a significantly higher activity in the Cue epoch of the PPM trials and Movement epoch of DM trials, likely reflecting initial planning processes in these epochs/conditions.

Another factor deals with the problem of dissociating visual target cue eccentricity from movement distance. As it may be argued, the more eccentric the visual cues, the more activity they can evoke, due to the relative over-representation of visual periphery in the parietal cortex (e.g.: Colby et al., 1988; Baizer et al., 1991; Motter and Mountcastle, 1981). In our study target eccentricity itself was inevitably correlated with the length of the visual trajectory of the end-effector (i.e. the placement of targets in the “big-step” conditions was more eccentric than in the “small-step” conditions). Therefore the increase in BOLD-signal that we observed in the early planning could have equally likely reflected any of these aspects. If this was true, however, that this effect should then be also visible in the cue phase of the DM trials, and, potentially, in the primary visual cortex. Both were clearly not the case. Hence the eccentricity of the visual target cue is unlikely to explain the observed results.

Movement duration can be considered yet another potential confounding factor. Movements towards more visually distant locations lead to longer lasting sensorimotor representations, which in turn may lead to higher BOLD activity (due to the long time constant of the BOLD-signal such change in motor duration will foremost surface as a change in signal amplitude). In our current study, however, this should again affect not only signal amplitudes during the movement epoch of the DM, but also those of the PPM. Moreover, duration-related signal changes should be visible also in primary motor cortex. Yet, such effect is lacking as well. Furthermore, movement durations cannot explain the gain effect we see in the CUE phase of the PPM trials. It thus seems plausible to conclude that the observed BOLD-signal modulation during the cue epoch in PPM and during the movement epoch in DM does solely reflect visual differences in the planned movement.

Finally, was the gain-related modulation of the BOLD-signal in DM related to movement planning or due to movement execution? The lack of gain-related BOLD signal modulation during the movement epoch of PPM trials suggests that the observed modulation in DM is rather related to planning differences between “big-step” and “small-step” conditions (present in DM) than to any immediate sensory or somatosensory feedback about the target or the actual movement (present both in DM and PPM).

For the above reasons we believe that that the gain-related modulation of BOLD-responses in PPC and PMd, occurring during the movement epoch of the direct movement condition and during the cue epoch of the pre-planning condition, is best explained by early planning processes, reflecting the visual consequences of upcoming movement.

### Visual action planning in PPC and PMd and its putative implications

The presence of a visual modulation of planning activity in human PPC and PMd is well in line with the known properties of both areas, as has been laid out in detail in the introduction. More generally, it supports the view that the visual movement consequences are a superordinated kinematic component of movement planning, determining the choice of appropriate dynamics in order to move the effector along the desired visual trajectory (Wolpert, 1995; Morasso, 1981).

The representation of visual consequences of a planned action appears more robust in PPC than in PMd. If we assume a processing hierarchy between these areas, our findings suggest that PPC delineates a rather general and abstract action plan in visual terms, which is subsequently translated into more specific motor programs by PMd (Desmurget and Sirigu, 2009; Kalaska and Crammond, 1995; Cisek and Kalaska, 2002; also compare to Westendorff et al., 2010). Yet we show that, contrary to what some of the above research may suggest, the sensory action representations are also present in PMd. This indicates that the both the posterior parietal parietal and premotor regions represent action plans based on action’s sensory outcome.

Such high-level visual representation of the outcome of an intended movement seems to be important for several aspects of action planning. When hand movements need to avoid obstacles and an appropriate trajectory has to be planned upfront it has to be done in visual terms. Moreover, simulating the desired visual outcome can serve as a stable reference for planning whenever effector efficiency is altered (i.e. due to fatigue or injury). This in turn requires an appropriate adaptation of motor programs that considers the current efficacy of the motor system and that is possibly realized via reciprocal cerebro-cerebellar connections with only limited involvement of awareness (Blakemore and Sirigu 2003).

Planning motor actions by simulating their visual consequences, is supposedly one of the vital prerequisites enabling effector selection and tool use. Actions engaging different end-effectors such as one’s bare hand or a stick obviously require different motor programs, even if the goals to be achieved are the same. The predictive representations of action outcomes allow for selecting optimal plans and evaluate them in advance, thus prevent acting on trial-and-error basis. Such evaluation allows also modifying the natural motor repertoire by incorporating the available end-effectors (e.g. tools, computer interfaces or even virtual environments) to achieve a desired sensory outcome. Such flexibility in planning would then broaden the spectrum of potentially available goals and actions (Gallivan et al., 2013; Haruno et al., 2001; Iriki et al., 1996; Maravita and Iriki, 2004) permitting a more efficient selection of both.

Finally, it could be further speculated that a representation of the visual consequences of planned actions in PPC and PMd also underlies our capacity to distinguish self- from externally- produced visual events (e.g. compare Synofzik et al. 2006). While this distinction has been mainly thought to be drawn from a comparison of an efference-copy based prediction of the visual consequences of self-action with the actual visual afference (Sommer and Wurtz, 2002), others suggest that this capacity may likewise refer to a comparison between *desired* and *actual* visual action outcomes (Bahcall and Kowler 1999; Synofzik et al. 2006).

Certainly, the exact role of PPC and PMd in these abovementioned functions remains to be determined. Yet, it is important to stress that a seemingly simple principle, i.e. the planning of action based on desired visual consequences, could have implications for a wide variety of functions extending beyond the motor domain.

## Conclusions

Our findings suggest that early planning activity in human posterior parietal cortex represents the visual consequences of planned actions independent of the actual motor programs required to realize these plans. Moreover, we found similar activity in human dorsal premotor cortex, suggesting that the two brain regions may collaborate in representing a visually defined action plans and, potentially, in translating them into appropriate motor commands. At this stage we may speculate that posterior parietal cortex, a region bridging between visual and motor areas might serve as the main driving force of this parieto-frontal planning system, utilizing information from bothie sources in order to create an effective movement plan.

## Acknowledgments

This work was supported by grants from BMBF (FKZ 01GQ1002) and DFG (CIN).

---

**Figure S1.**
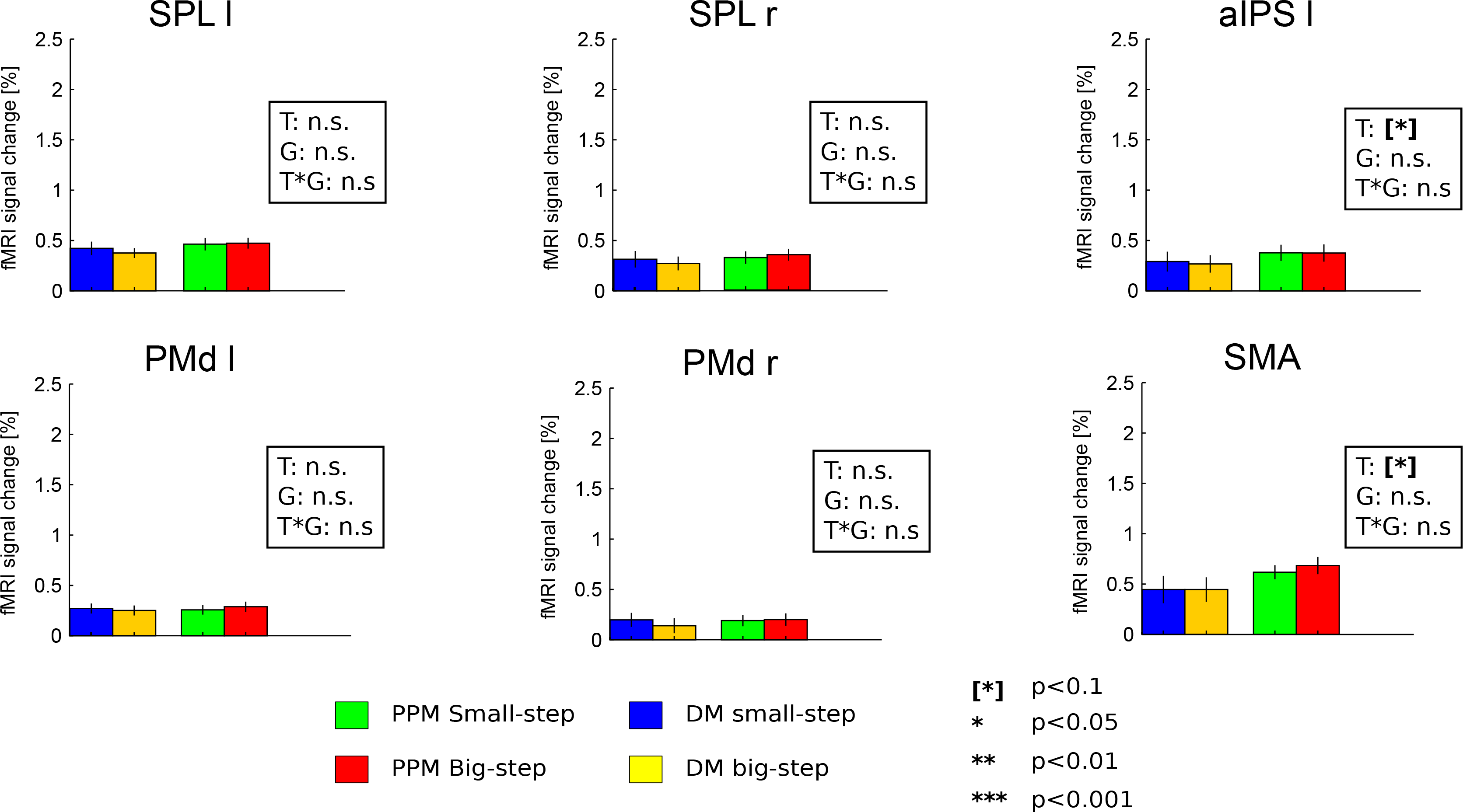
BOLD activity in all ROIs during the delay epoch. Sustained activity was present in all ROIs. Signal increases in PPM with respect to DM in M1 could relate to unspecific motor preparation. All values represent averages calculated across subjects’ mean activity +/− SEM.

**Figure S2.**
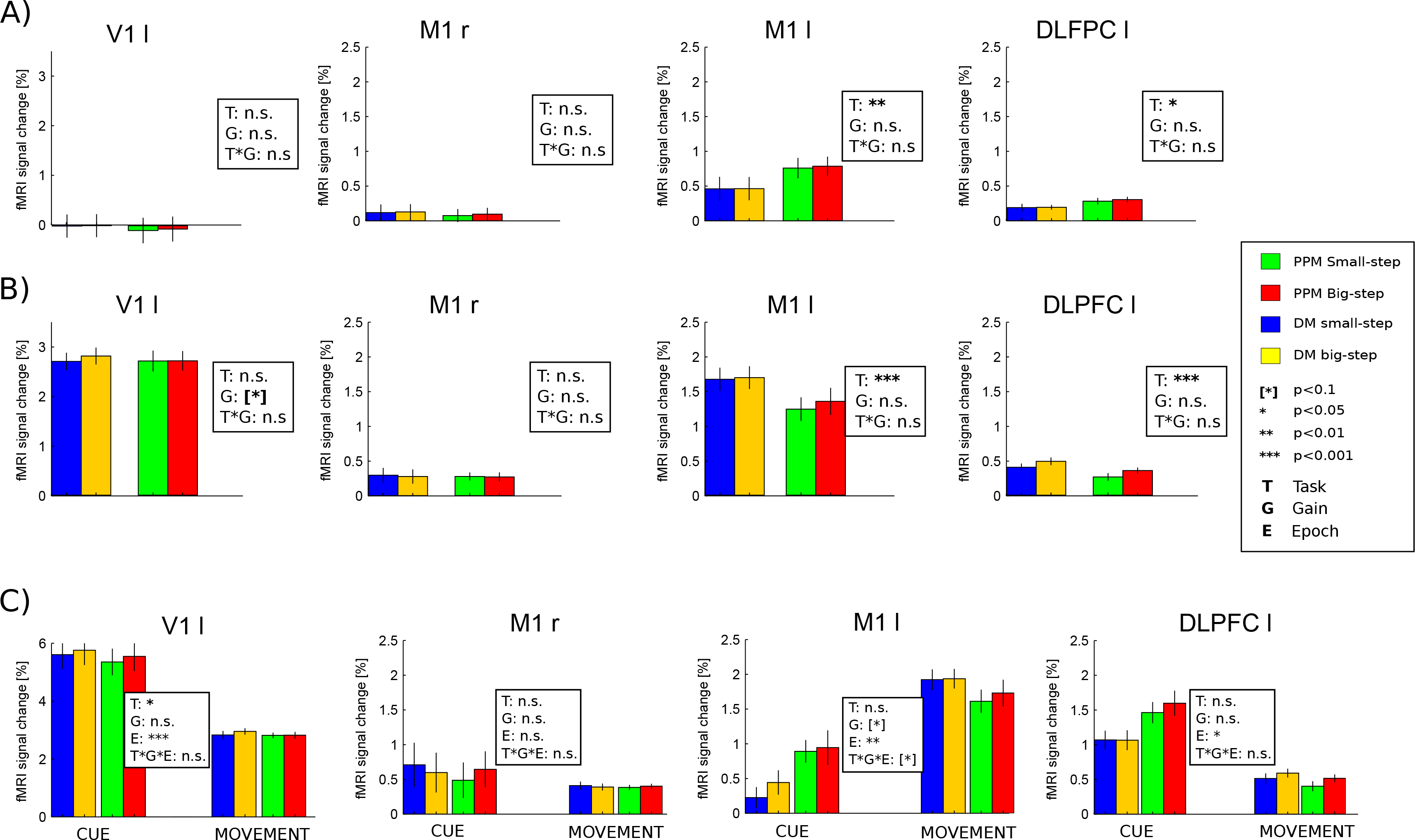
Delay (A) and movement epoch (B) betas extracted from primary visual and right primary motor areas. The V1 activity shows a weak trend related to movement gain in the movement epoch of DM trials. C) Control ROI betas extracted in the cue and movement phases, used to capture early planning activity. All values represent averages calculated across subjects’ mean activity +/− SEM.

